# Coevolution-informed Bayesian optimization for sample-efficient protein design

**DOI:** 10.64898/2026.08.06.743295

**Authors:** Dulitha P. Kulathunga, Divyanshu Shukla, Davit Potoyan

**Affiliations:** Department of Chemistry, Iowa State University, Ames, IA 50011; Bioinformatics and Computational Biology Program, Iowa State University; Roy J. Carver Department of Biochemistry, Biophysics and Molecular Biology, Iowa State University, Ames, IA 50011

## Abstract

Protein engineering is limited less by generating variants than by the cost of evaluating them, so designing under a tight budget demands sequence features that let a model learn fitness from very few examples. We introduce ALSEBO (Active Learning Sequence Exploration via Bayesian Optimization), which couples a generative latent sequence landscape to Bayesian optimization and featurizes candidates with direct-coupling-analysis (DCA) coevolutionary statistics. This representation carries a specific inductive bias: it places the dominant organizer of the fitness landscape along a single linear coordinate, producing a smooth, funnel-like objective that a low-data surrogate navigates efficiently. On a virtual avGFP fluorescence benchmark, ALSEBO reaches the optimum in ∼40 evaluations and outpaces protein-language-model embeddings and raw latent coordinates; controls with representation-neutral oracles confirm that the advantage is intrinsic, not an artifact of the benchmark. Molecular dynamics of the optimized variant recovers structural hallmarks of fluorescence, and ALSEBO transfers to divergent GFP orthologs and to a non-GFP enzyme, establishing a data-efficient route to protein design.

## 1 Introduction

Protein engineering faces a fundamental scaling problem: a sequence of modest length *L* occupies a combinatorial space of size 20*^L^*, of which only a vanishingly small fraction folds into stable, functional structures. ^1–4^ Evolution has navigated this space by iterative selection under biophysical constraints over geological timescales; ^5,6^ modern engineers cannot. Rational design is limited by incomplete knowledge of sequence–structure–function relationships and rarely captures long-range epistasis; ^7,8^ directed evolution treats the protein as a black box and rarely crosses deep fitness valleys; ^9^ semi-rational saturation mutagenesis encounters combinatorial explosion at even modest mutation counts.^10–12^ Under realistic experimental budgets, the common bottleneck is the same: exploration footprints are infinitesimally small relative to the accessible sequence space. ^13–15^

Machine-learning approaches have partially addressed this limitation through two complementary ideas. Generative models, including Variational Autoencoders (VAEs),^16^ Protein Language Models (pLMs), ^17^ and diffusion models, ^18^ learn the grammar of protein sequences and compress it into structured latent manifolds enriched for viable variants; structure-based generators such as ProteinMPNN and RFdiffusion additionally design sequences and backbones de novo. ^19,20^ Bayesian Optimization (BO) paired with active learning then converts fitness prediction into a decision-making problem by balancing exploitation of known peaks against exploration of uncertain regions.^21–23^ Each idea addresses a piece of the problem but not the whole: generative models produce biophysically plausible sequences without optimizing function, while BO formulations on raw discrete sequence spaces ^24^ or on naive latent coordinates ^25^ either suffer the curse of dimensionality or fall into decoder cliffs where latent steps fail to produce viable folds. Closing the gap requires a candidate space with a built-in bias toward structural plausibility and a feature representation that lets a low-N surrogate learn fitness quickly.

Here we introduce ALSEBO (Active Learning Sequence Exploration via Bayesian Optimization), a framework that couples a generative latent sequence landscape ^26,27^ with a Gaussian-process surrogate and active learning to find high-fitness variants from very few evaluations, and that works with any candidate generator and objective. Its central design choice is the sequence representation: ALSEBO featurizes candidates with Direct Coupling Analysis (DCA), the coevolutionary statistics of a protein family. This choice carries a specific inductive bias: an algebraic identity places the DCA coevolutionary energy along a single coordinate of the feature vector, making the dominant organizer of the fitness landscape directly accessible to a low-data surrogate. Benchmarking DCA features against protein-language-model (ESM-2) embeddings ^17^ and raw latent coordinates on a virtual avGFP fitness landscape,^28^ we find that DCA reaches the optimum in the fewest evaluations, traceable to a smoother, funnel-like objective surface. Because the avGFP oracle and the optimizer share the DCA representation, we retest the advantage against representation-neutral oracles that carry no coevolutionary information, and against a non-coevolutionary target (the enzyme Ube4b), confirming that it is intrinsic rather than an artifact of the benchmark. ALSEBO further transfers to the divergent GFP orthologs cgreGFP and amacGFP,^29^ establishing a data-efficient route to protein engineering under realistic experimental budgets.

## 2 Results

ALSEBO proceeds in four stages (Figure 1): it draws a candidate library from a latent generative landscape, featurizes each sequence, initializes a diverse training set, and then runs an activelearning loop in which a Gaussian-process surrogate and an upper-confidence-bound acquisition function choose the next batch to evaluate. The framework is agnostic to the sequence generator, feature representation, and objective function; full details are given in Methods.

**Figure 1:**
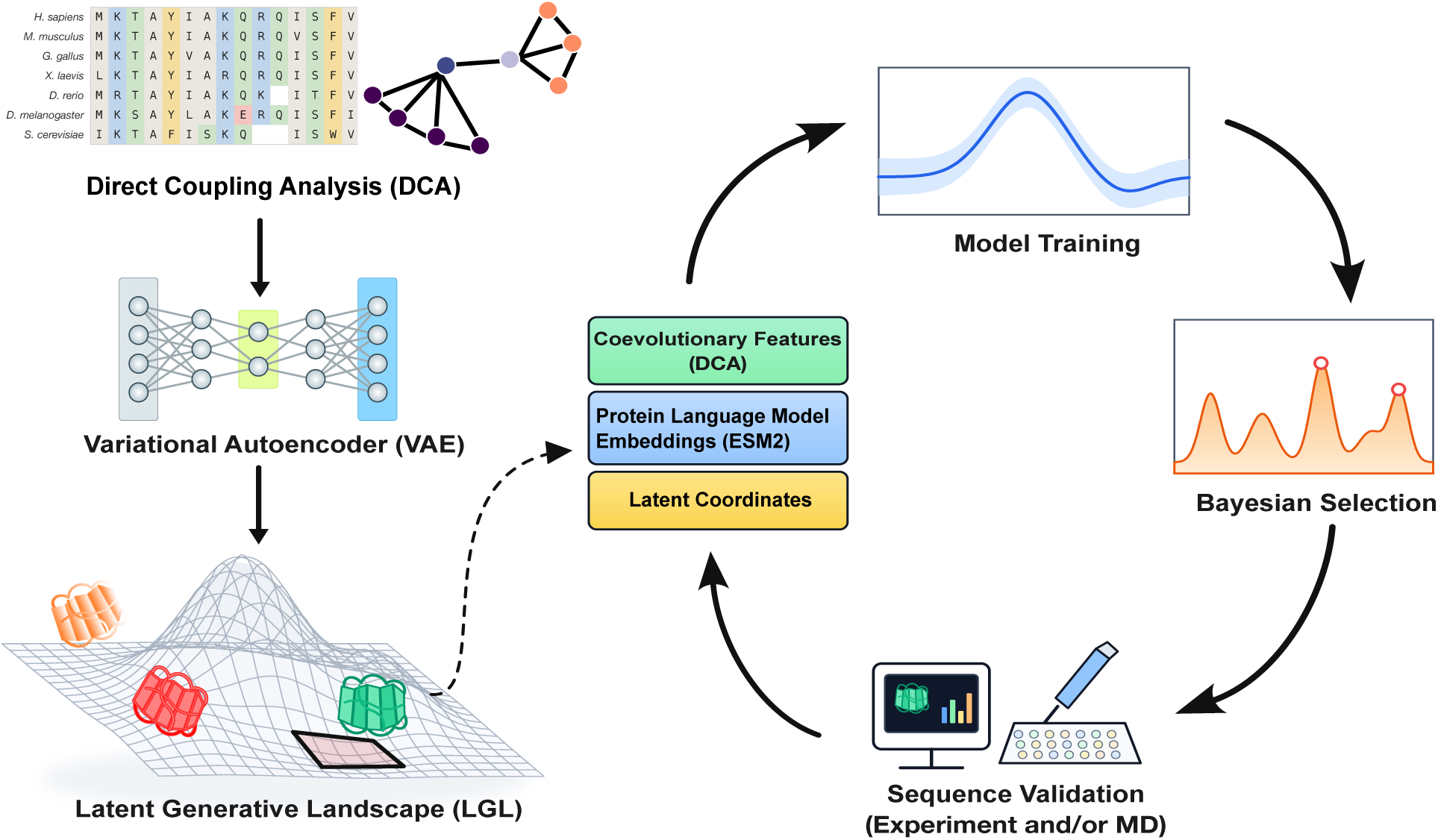
ALSEBO workflow. (A) A VAE trained on a multiple-sequence alignment embeds sequences in a two-dimensional latent space; a DCA model trained on the same alignment scores decoded sequences across the latent grid to form the latent generative landscape (LGL), coloured by DCA Hamiltonian. (B) Candidate sequences are featurized (coevolutionary, language-model, or latent features) and passed to an active-learning loop in which a Gaussian-process surrogate and Bayesian optimization prioritize the next sequences for evaluation, with results fed back to update the surrogate.

We constructed the latent generative landscape on a 500 × 500 grid over (*z*_0_*, z*_1_), decoding the MAP sequence at each grid point and scoring it with the DCA Hamiltonian (Figure 1). The resulting scalar field is structured into low-Hamiltonian basins separated by higher-energy barriers, with known family members embedded in the low-energy regions; this is the candidate space for ALSEBO’s active-learning loop (Figure 1).

For the avGFP case study, the GFP-family landscape organizes into low-Hamiltonian basins that partition sequences into phenotypically related clusters (Figure 2A). Centered on the basin containing the wild-type embedding, we sampled the surrounding low-energy region densely and decoded 2,143 candidate variants (Figure 2B). The resulting library compresses the nominally intractable 20*^L^* sequence space into a tractable set while preserving substantial diversity; mutation counts range from 0 to 60 relative to the wild type (Figure 2E), indicating that latent-space sampling guided by the DCA Hamiltonian restricts the search to structurally viable regions without collapsing onto near-wild-type variants.

**Figure 2:**
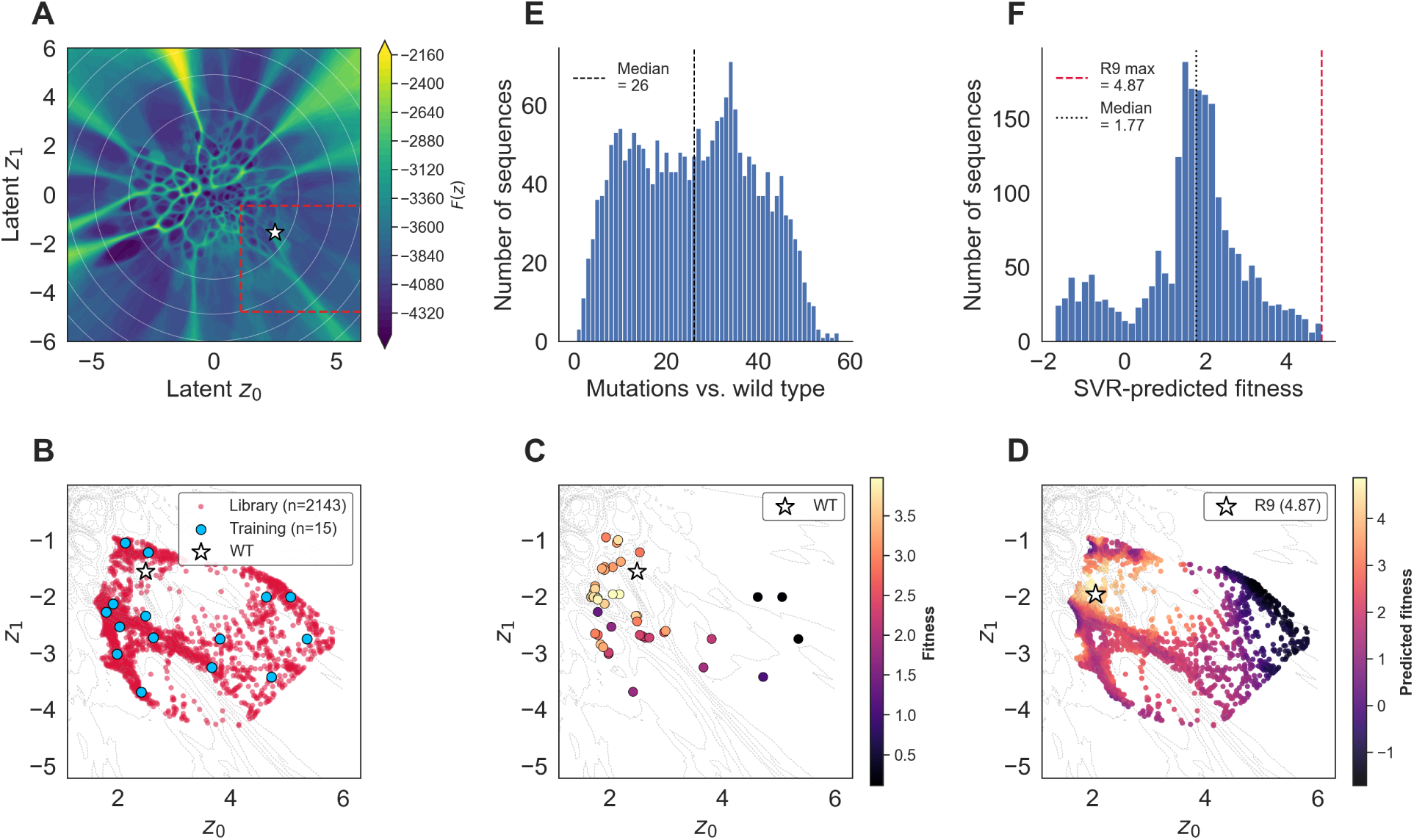
GFP latent generative landscape and candidate library. (A) Effective-score landscape *F* (*z*) = − log *π_θ_*(*z*) + ⟨*H*_DCA_⟩ (Eq. 9) over the VAE latent domain [−6, 6]^2^; white contours mark prior iso-levels, the white star is wild-type GFP, and the red dashed rectangle is the library bounding box (panels B–D). (E) Distribution of mutation counts in the 2,143-sequence library relative to wild type (0–60 substitutions, median 26). (F) Distribution of SVR-predicted fitness across the library; black dotted line, median; red dashed line, the R9 maximum. (B–D) Zoomed views on the library footprint, sharing axes, with a thin grey Hamiltonian contour for orientation: (B) library sequences (crimson) and the initial *k*-means-centroid training set (blue); (C) BO-evaluated sequences coloured by SVR-predicted fitness; (D) all library sequences coloured by SVR-predicted fitness, R9 marked (white star).

### 2.1 Establishing the Virtual Fitness Landscape for avGFP

Applying the virtual objective to the 2,143-sequence generated library yields a multimodal fitness distribution (Figure 2F) whose maximum falls at predicted fitness 4.87 on a nine-mutation variant we designate R9: Y37N, S63T, S70A, F97S, N103T, Y143F, M151T, V161A, I169V. Projecting the oracle scores back onto the DCA manifold (Figure 2D) visualizes the search landscape the optimizer navigates. Eight of R9’s substitutions (with the two-residue alignment offset: Y39N, S65T, F99S, N105T, Y145F, M153T, V163A, I171V) coincide with the hallmark consensus mutations of superfolder GFP.^30^ This biological interpretability of R9 makes it a meaningful anchor for the active-learning and MD analyses that follow.

### 2.2 Navigating Fitness Landscapes via DCA-Informed Bayesian Optimization

We ran three independent BO campaigns targeting R9 (the oracle’s maximum over the generated library), initialized with *T* = 5, 10, or 15 diverse starting sequences to probe sensitivity to the initial prior. Generated sequences were featurized via DCA encoding (Eq. 10) before GPR surrogate training.

Across all three initial budgets, ALSEBO located the oracle optimum (R9) within 6–8 active-learning cycles, corresponding to 40–45 total sequence evaluations (Figure 3C). The trajectories projected onto the DCA-feature PC plane (Figure 3A) and onto the VAE latent space (Figure 3B) are not random walks: early cycles explore broadly, and successive rounds converge into the basin containing R9. Panel 3C makes the exploration-exploitation balance concrete: the best sequence per batch rises monotonically, while periodic dips in the batch mean are consistent with the UCB acquisition function sampling in regions of high predictive variance, preventing premature collapse onto local peaks. The temporal evolution of *σ*(*x*) and *a*(*x*) (SI Figure 1) is consistent with this picture, showing a transition from broad exploration at *t*= 1 to targeted exploitation of the identified high-fitness basin by *t* = 7.

**Figure 3:**
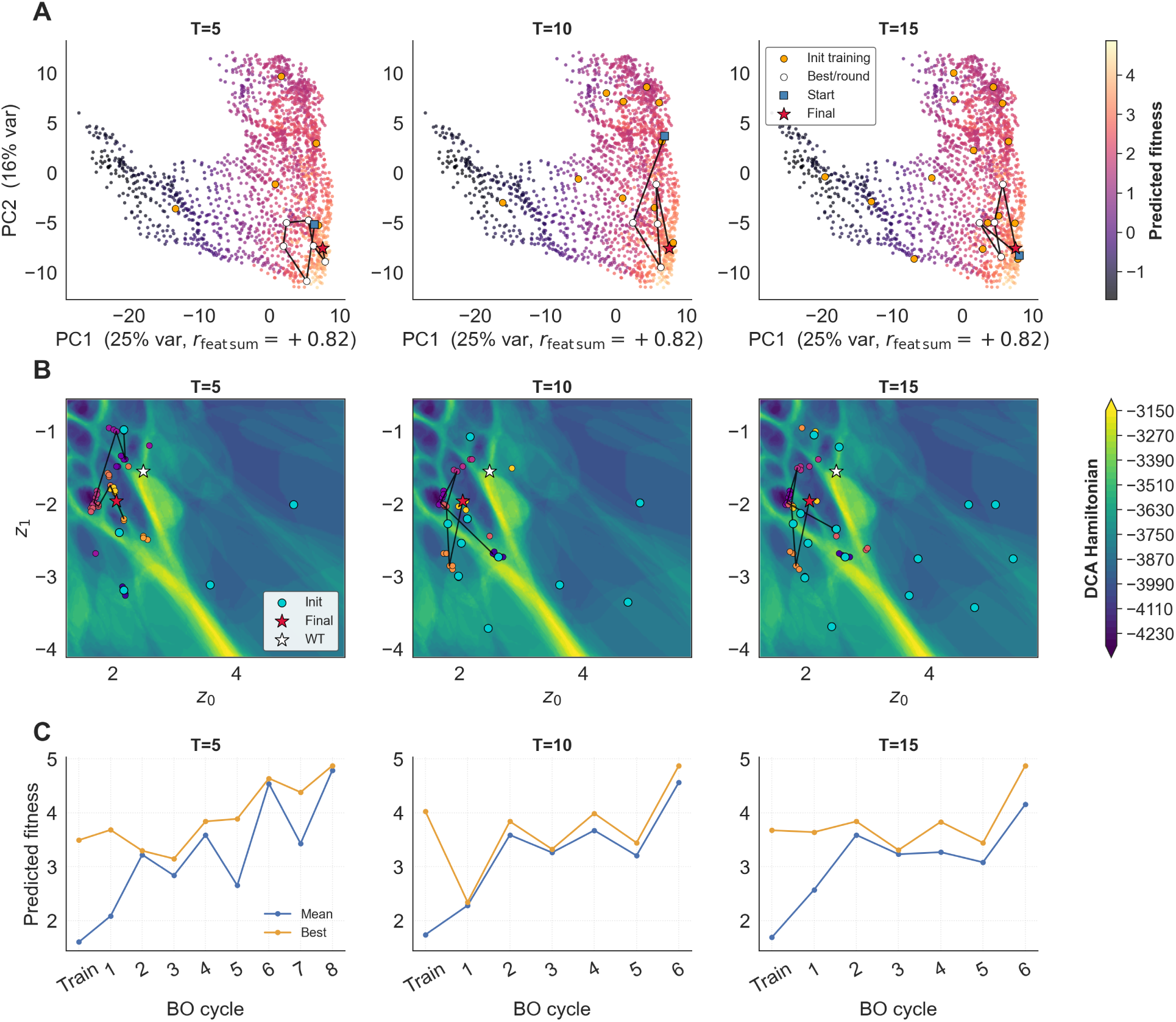
ALSEBO campaigns under constrained initial data. Columns: DCA-featurized campaigns initialized with *T* = 5, 10, or 15 starting sequences (left to right). (A) Optimization paths on the first two principal components of the DCA feature space, coloured by predicted fitness; PC1 correlates *r* = 0.82 with the feature-sum direction (Eq. 11) and *r* = 0.89 with SVR fitness. Orange disks, initial training sequences; black line, best-scoring sequence per round; red star, final optimum. (B) The same trajectories on the LGL Hamiltonian landscape in (*z*_0_*, z*_1_): every evaluated sequence (cycle-coloured discs), initial training sequences (cyan), best-per-round path (black line), wild type (white star), and final optimum (red star). (C) Cycle-wise mean and best fitness within each BO batch.

The trajectories in Figure 3B stay confined to the low-Hamiltonian basins of the LGL, consistent with the DCA-featurized surrogate inheriting the generative prior’s structural-stability constraint, even though the oracle never penalizes Hamiltonian directly. At the same time, the converged solutions depart substantially from natural sequence variation (R9 carries nine epistatically coupled substitutions not found together in the training alignment), suggesting that the DCA inductive bias acts as a stability regularizer rather than a phylogenetic trap. Furthermore, SI Figure 4 highlights how per-position mutational frequencies increase towards optimal sequence mutations in each stage of the active learning cycle.

### 2.3 Feature Representation and Landscape Topography Govern Optimization Efficiency

Surrogate performance in low-data regimes depends on the inductive bias carried by the feature representation. We compare three information-dense alternatives to one-hot encoding (which is additive and, by construction, ignores epistasis) on our in silico oracle: 235-dimensional DCA features, 640-dimensional ESM-2 embeddings, and the raw 2D LGL latent coordinates. Across 100 independent campaigns the three representations separate cleanly: DCA-featurized optimization converges to the oracle optimum fastest, ESM-2 next, and raw latent coordinates slowest (Figure 4A,B). With random initialization (Figure 4C), DCA and ESM-2 both reach a 1.00 success rate, versus 0.95 for latent coordinates and 0.03 for random search, while mean steps to optimum separate cleanly: 31.1 ± 9.5 (DCA), 36.3 ± 8.1 (ESM-2), and 59.7 ± 17.6 (latent).

**Figure 4:**
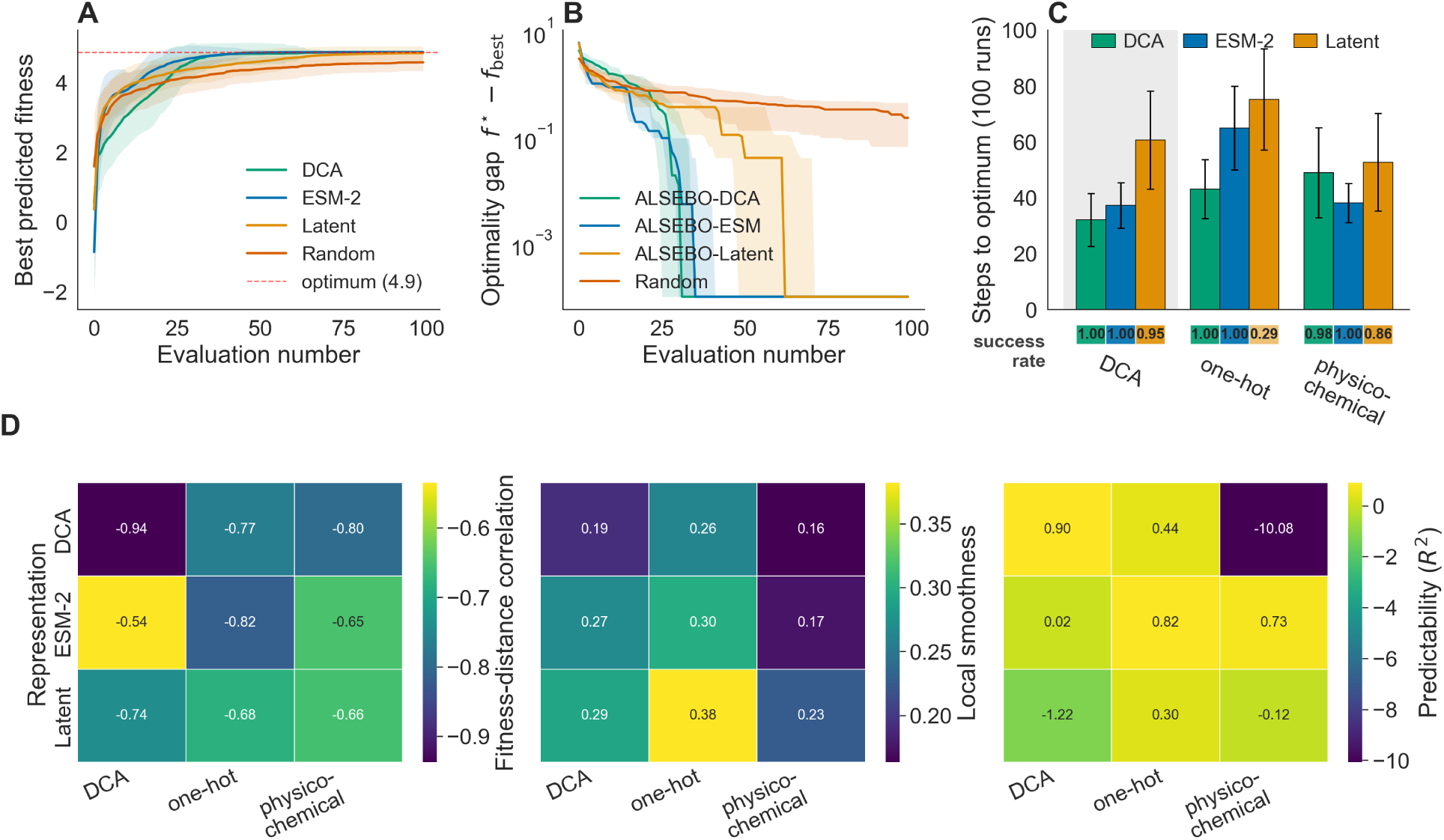
Feature-representation benchmark and landscape-topology controls. (A) Best predicted fitness versus number of sequences evaluated (mean over 100 independent campaigns; shaded band, SD) for DCA, ESM-2, and latent features against a random-search baseline; the red dashed line marks the oracle optimum (≈ 4.9). (B) Optimality gap *f* ^∗^ − *f*_best_ on a log scale versus evaluation number (mean over 100 campaigns; shaded band, SD) for the same methods. (C) Mean steps to reach the optimum over 100 random initialized campaigns for each feature representation (DCA, green; ESM-2, blue; latent, orange; error bars, SD) under three oracles trained on the same fluorescence data: the DCA-based oracle (grey shaded) and the representation-neutral onehot and physicochemical oracles. The success rate (fraction of campaigns reaching the optimum) is annotated beneath each bar. (D) Landscape-topology characterization of each feature representation (rows: DCA, ESM-2, latent) under each oracle (columns: DCA, one-hot, physicochemical): Fitness–Distance Correlation (left; more negative indicates a stronger global funnel), local smoothness (middle), and GP predictability *R*^2^ (right).

To understand this hierarchy, we characterized the topology of the objective landscape in each feature space (Figure 4D, SI Figure 7). DCA’s sample efficiency reflects a funnel-like landscape with a strong global gradient (Fitness-Distance Correlation, FDC = −0.94) that guides the GP surrogate toward the optimum. The 2D latent space preserves a macroscopic funnel (FDC = −0.74), but its extreme compression leaves too little local resolution for the surrogate, giving poor predictability (*R*^2^ = −1.22) and a large evaluation budget. ESM-2 yields a more rugged, dispersed topology (FDC = −0.54), yet its high-dimensional embeddings retain enough local predictability to navigate effectively. By encoding pairwise epistasis, DCA smooths the mutational cliffs that confound optimizers, letting the GP exploit high-fitness basins directly. This topological view is complementary to the algebraic one: the feature-sum identity (Eq. 11) confines the fitness-organizing signal to a few dimensions of the DCA space, which is geometrically the funnel of Figure 4D and, information-theoretically, means each evaluation is highly informative, so the GP-UCB regret bound reproduces the observed sample-efficiency ordering (SI Figure 6).

Because the avGFP oracle is a DCA-SVM (chosen for its high accuracy on the experimental data, and so the most realistic surrogate), it shares a representation with the DCA-featurized optimizer, which could inflate DCA’s advantage. To separate this shared-basis effect from genuine optimization power, we rebuilt the oracle from one-hot encodings and from physicochemical descriptors, neither of which encodes coevolution, and repeated the benchmark against each (Figure 4C). The latent representation now collapses (success rate 0.29 on the one-hot landscape), confirming that a 2D space over-smooths the high-frequency epistasis needed to pinpoint the optimum.

ESM-2 proves a robust generalist: with no shared basis with the oracle, it forms strong funnels for both the one-hot (FDC = −0.82) and physicochemical (FDC = −0.65) objectives, keeps a 1.00 success rate, and even outpaces DCA on the physicochemical landscape (38.4±7.2 steps). Critically, DCA holds up as well: without a shared representation, it retains a 1.00 success rate on the onehot landscape and 0.98 on the physicochemical one, supported by strong global gradients (FDC = −0.77 and −0.80). Its advantage is therefore not memorization of the oracle’s architecture but a transferable, biologically grounded coordinate system that clusters functional variants.

### 2.4 Structural Validation and MD Simulation Analysis

High surrogate-predicted fitness does not guarantee a structurally viable protein: computationally optimized variants can still misfold or destabilize in solution. To test whether ALSEBO’s predictions correspond to architectures consistent with fluorescence competence, we ran 1 *µ*s allatom MD simulations on five variants spanning the optimization trajectory (R6, R7, R8, R9, R10) alongside the S65T baseline (W2). The R9 mutations are distributed across the *β*-barrel scaffold (Figure 5A); the remaining variants carry overlapping but distinct subsets, including the destabilizing L62F substitution in R10. We emphasize that MD simulates *structural correlates* of fluorescence (backbone rigidity, chromophore hydration, chromophore geometry), not the emission observables themselves; quantum yield, excited-state relaxation, and chromophore maturation are outside the scope of ground-state classical MD and would require either quantum-chemical treatment of the excited state or direct in vitro characterization. The simulations per variant consist of single 1 *µ*s trajectories and are therefore best read as qualitative mechanistic support rather than as quantitatively tight estimates of subtle differences. We evaluate the simulations against the well-established photophysical logic of green fluorescent proteins: fluorescence requires a rigid *β*-barrel that shields the chromophore from solvent, and a planar chromophore geometry preserving *π*-conjugation between the imidazolinone and phenolate rings. ^31^ Each MD observable below probes one step in this causal chain.

**Figure 5:**
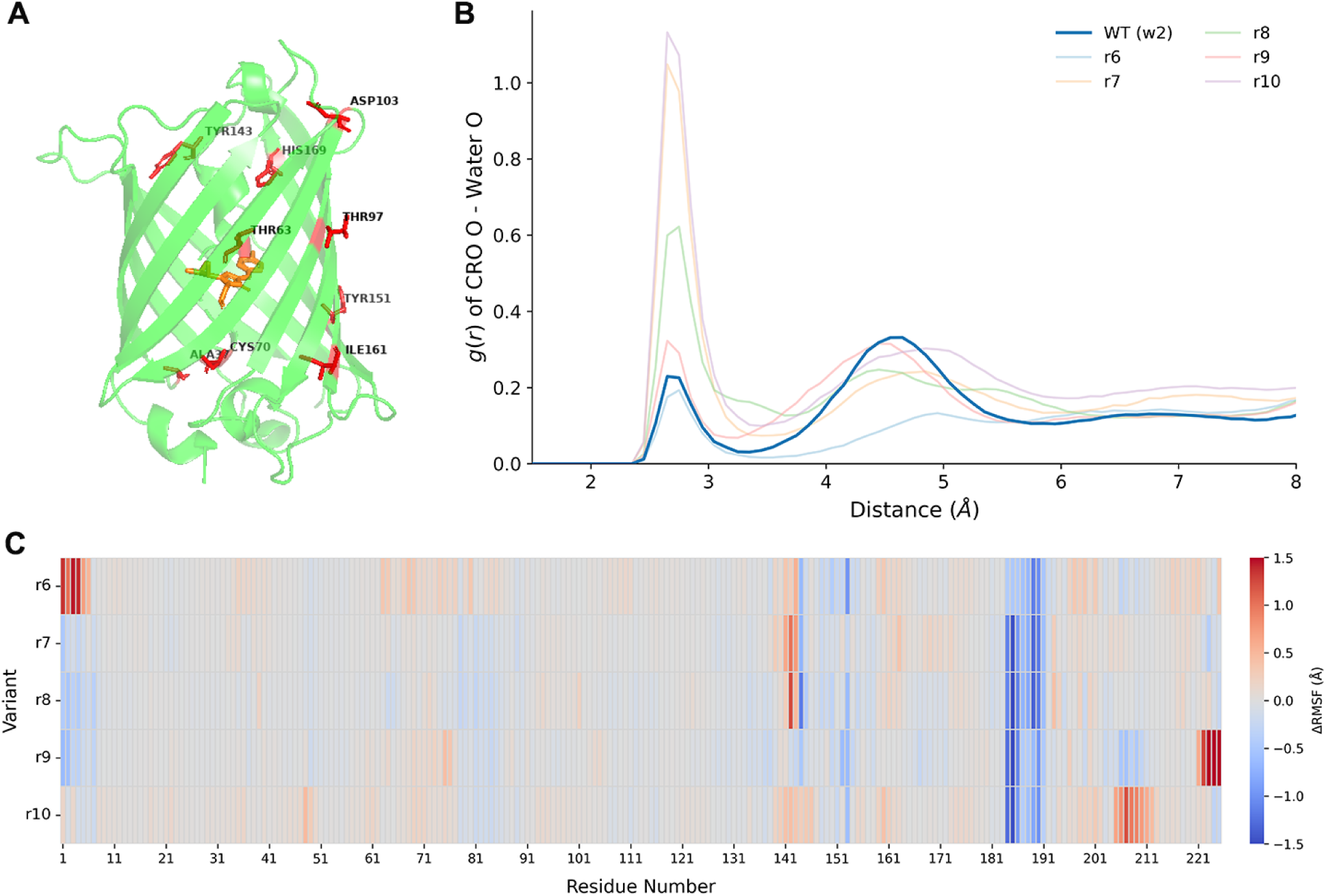
Molecular-dynamics analysis of ALSEBO-optimized variants. (A) The nine R9 mutations mapped on the avGFP *β*-barrel scaffold. (B) Radial distribution function *g*(*r*) of water oxygen atoms relative to the chromophore phenolate oxygen, for W2 and R6–R10. (C) ΔRMSF heatmap of R6–R10 relative to the W2 (S65T) baseline, by residue.

**Step 1: barrel rigidification.** Backbone C*α* RMSF profiles (**SI Figure 8A**) show that R9 reproduces and slightly sharpens the dynamic signature of the W2 baseline despite carrying nine sub-stitutions, while the destabilized R10 variant develops flexibility spikes exceeding 1.0 Å at residues 140–150 (gatekeeper loops) and 205–215. The ΔRMSF heatmap relative to W2 (Figure 5C) resolves this further: all optimized variants show a consistent ΔRMSF *<* −1.0 Å along the *β*-strand 9-10 seam (residues 180–195), indicating that the superfolder-like mutational core synergistically rigidifies the barrel’s load-bearing strands. Intermediate variants reveal the optimization trajectory in structural terms: R6 is indistinguishable from W2 (early, conservative exploration), R7 and R8 show emergent rigidification paired with localized gatekeeper destabilization, and R9 is the fitness node where global compaction is achieved without sacrificing active-site anchors.

**Step 2: hydration exclusion.** The rigidified barrel translates directly into macroscopic compaction: R9 exhibits a narrower SASA distribution (∼11,250 Å^2^) centered on a lower mean than W2 (∼11,350 Å^2^, SI Figure 3), whereas R7 and R10 populate broader distributions with tails at high surface area, the structural correlate of their gatekeeper loop defects. This compaction is what determines chromophore solvation. Radial distribution functions *g*(*r*) of water oxygens relative to the chromophore phenolate oxygen (Figure 5B) show that R9 maintains a near-desolvated primary hydration shell (coordination number CN = 0.32), while R7 and R10 develop prominent first-shell peaks with CN = 1.04 and 1.14, indicating one water molecule directly hydrogen-bonded to the phenolate. Because water is an efficient vibrational sink, direct coordination provides a non-radiative decay pathway that extinguishes fluorescence; R9’s desolvated geometry is precisely what is required to avoid this quenching channel.

**Step 3: chromophore planarity.** Within the compact barrel, R9 preserves the packing contacts that position the chromophore (SI Figure 8B): near-unity contact frequencies with VAL146, THR199, and PHE161, together with a strong 0.77 packing frequency at position 141 despite the Y→F mutation at that site (a hallmark sfGFP substitution that relieves local steric clash). The suboptimal variants fail this test in specific ways: R6 loses PHE161 contact (0.71) and forms an atypical HIE177 interaction absent in both R9 and W2; R10’s L62F substitution disrupts the chromophore pocket, reducing the PHE141 contact to 0.34. The consequence of preserved packing is that the imidazolinone and phenolate rings of the chromophore remain coplanar across all optimized variants (SI Figure 2): free-energy basins are confined to *ϕ* ≈ 0^◦^, *τ* ≈ ±180^◦^ with no accessible populations at the twisted, non-fluorescent ±90^◦^ configurations.

Taken together, the simulations trace a structurally coherent chain of correlates from sequence to photophysics: the R9 mutational pattern is associated with barrel rigidification, the rigidified barrel with reduced water density near the chromophore, and the preserved packing cavity with chromophore planarity. R10 departs from this chain at the first step (L62F-induced destabilization), with downstream consequences in water coordination and packing. Sub-optimal variants (R6, R7, R8) show partial failures at individual links. That the DCA-featurized GPR ranks these variants consistently with three independent biophysical scales (backbone fluctuation, hydration, chromophore geometry) indicates the surrogate has learned mechanistically interpretable correlates of fitness. These are structural correlates, not a measurement of fluorescence; replicate trajectories and direct photophysical or experimental readouts are the natural next step.

### 2.5 Generalization across Protein Families and Design Objectives

To further evaluate the generalizability of ALSEBO beyond the original avGFP benchmark, we constructed two additional GFP fitness landscapes based on the cgreGFP and amacGFP orthologs reported by Gonzalez Somermeyer et al. ^29^ These systems represent varying evolutionary distances from avGFP, with amacGFP (*Aequorea macrodactyla*, Hydrozoa) sharing approximately 82% sequence similarity and cgreGFP (*Clytia gregaria*, Hydrozoa) sharing approximately 41% sequence similarity. ^29^ Following the same strategy used for the avGFP benchmark, experimentally charac-terized mutational datasets from Gonzalez Somermeyer et al. were used to train SVM models over DCA-derived features, which served as virtual objective functions for fluorescence fitness prediction. Homologous sequence datasets were then collected through multiple sequence alignments and used to generate latent fitness landscapes using the LGL-VAE framework. For cgreGFP and amacGFP we deliberately built more diverse libraries than for avGFP, including insertions and deletions relative to the reference, which produced more rugged landscapes that stress-test the optimizer. The two systems pose complementary challenges: amacGFP forms one broad continuous basin with narrow barriers, whereas cgreGFP was sampled from two high-fitness basins separated by an energy barrier. Both are harder to navigate than the smooth, localized avGFP landscape. The diverse nature of these generated landscapes is quantitatively supported by their mutational profiles (SI Figure 9), which demonstrate the broad mutational spread and complex, multi-modal fitness distributions required to rigorously stress-test the active learning framework. From only five initial sequences, ALSEBO still reached the high-fitness basins in both systems: 5 cycles (30 evaluations) for amacGFP and 9 cycles (50 evaluations) for cgreGFP, the larger budget reflecting cgreGFP’s barrier-separated topology. In both, the best and mean predicted fitness rose steadily across cycles, showing that the latent library and DCA features guide exploration even on rugged, heterogeneous landscapes.

We next tested ALSEBO on a non-GFP target with an entirely different objective. As a representative system we used ubiquitination factor E4B (Ube4b), for which large-scale deep mutational scanning data are available. ^32^ As the virtual objective we adopted the sequence–function convolutional neural network of Gelman et al. ^33^, trained on 98,297 Ube4b variants assayed for ubiquitin-ligase activity (Pearson correlation 0.68 between predicted and held-out experimental fitness). Importantly, this oracle is a CNN rather than a DCA-trained SVR, so here the DCA-featurized optimizer and the objective do not share a representational basis.

Using the same latent landscape-generation strategy, we sampled ∼4000 high-fitness Ube4b sequences of 102 residues in length, with a maximum of 2 gaps relative to the reference sequence, and evaluated them using the trained CNN model (Figure 6A). Bayesian optimization over this objective landscape identified an optimized sequence within 6 cycles, corresponding to only 35 total sequence evaluations. Notably, the optimized sequence shared only ∼20% sequence similarity with the initial Ube4b wild-type sequence, indicating that ALSEBO was able to explore a distant region of generated sequence space while maintaining high predicted activity.

**Figure 6:**
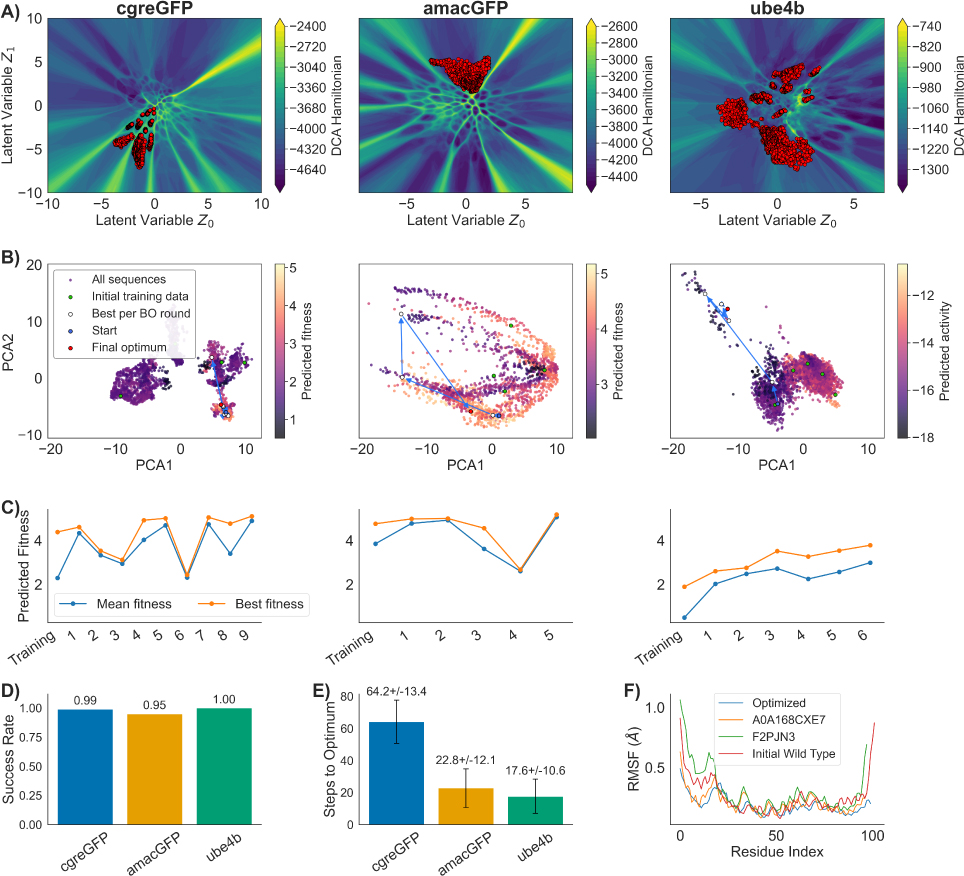
Generalization of ALSEBO across protein families and objectives. Columns correspond to the three test systems: cgreGFP, amacGFP, and Ube4b (left to right). (A) Generated latent fitness landscapes, with decoded candidates overlaid and coloured by each system’s virtual objective. (B) Bayesian-optimization trajectories visualized by t-SNE of the feature space, showing the initial training set, the best-per-cycle path, and the start and final-optimum points. (C) Per-cycle mean and best predicted fitness. (D) Success rate and (E) mean steps to the predicted optimum across 100 independent campaigns per system (cgreGFP, 0.99 / 64.2 ± 13.4; amacGFP, 0.95 / 22.8 ± 12.1; Ube4b, 1.00 / 17.6 ± 10.6). (F) Backbone C*α* RMSF of the optimized Ube4b sequence, its two nearest natural neighbours in latent space (UniProt F2PJN3 and A0A168CXE7), and the wild type.

To evaluate whether this highly divergent optimized sequence retained dynamic features consistent with functional Ube4b-like proteins, we performed 1.2 *µ*s all-atom molecular dynamics simulations of the initial wild-type sequence, the optimized sequence, and two natural sequences nearest to the optimized variant in the VAE latent space, UniProt IDs: F2PJN3 and A0A168CXE7. The main differences in RMSF (Figure 6F) were observed in the extended N-terminal region, which has been reported to exhibit auto-ubiquitination activity in vitro. ^32^ To assess structural consistency, we aligned the generated optimized structure with representative clustered structures obtained from the wild-type dynamics. The U-box domain of the optimized sequence aligned well with the initial wild-type structure, whereas larger alignment differences were localized to the extended N-terminal region. Together, these results suggest that the optimized sequence preserves Ube4b-like conformational dynamics and structural features of the functional U-box domain while occupying a distant region of sequence space.

To assess robustness, we performed 100 independent optimization campaigns with random initializations for all three systems. ALSEBO achieved convergence rates of 99%, 95%, and 100% for cgreGFP, amacGFP, and Ube4b, respectively. On average, the predicted global optimum was identified after 64.2 ± 13.4 sequence evaluations for cgreGFP, 22.8 ± 12.1 sequence evaluations for amacGFP, and 17.6 ± 10.6 sequence evaluations for Ube4b. The higher evaluation budget required for cgreGFP is consistent with its greater optimization complexity relative to both Ube4b and amacGFP: compared with Ube4b, cgreGFP is approximately three times longer in sequence length, increasing the dimensionality of the sequence–fitness relationship, while compared with amacGFP, the cgreGFP candidates were sampled from two high-fitness latent regions separated by an energy barrier, reflecting greater sequence diversity than the single-region amacGFP landscape.

Together, the GFP and Ube4b results show that ALSEBO transfers across protein families and objective types: given any objective, experimental or computational, it navigates large sequence landscapes to recover high-scoring candidates. This positions ALSEBO as an actionable pipeline for protein design beyond GFP-like systems.

## 3 Discussion

ALSEBO couples generative sequence modeling with coevolution-informed active learning to make protein optimization data-efficient, reaching high-fitness variants in tens of evaluations rather than the large screening libraries that directed evolution and deep mutational scanning require.

Two analytic properties explain why the framework is so sample-efficient. First, the latent generative landscape can be read as an effective score (Eq. 9) that combines two roles: the VAE prior acts as a soft domain regularizer, marking where the decoder produces trustworthy sequences, while the DCA Hamiltonian ranks sequences within that region. The two terms are not thermodynamically commensurate, so the result is a composite score rather than a true free energy, but it cleanly separates *where* to search from *how* to rank. Second, the DCA featurization satisfies an exact algebraic identity (Eq. 11): the coevolutionary Hamiltonian appears as a single linear direction of the feature vector. This is an inherent property of the representation, not a tuning choice, and it concentrates the dominant organizer of the fitness landscape into very few effective dimensions. The practical consequence, borne out by our landscape characterization, is a smooth, funnel-like objective surface (high FDC) that a low-data GP surrogate can navigate from only a handful of measurements. Crucially, this advantage is not an artifact of how the benchmark is scored: it persists when the oracle is rebuilt from unrelated sequence descriptions, one-hot encodings and physicochemical residue properties that carry no coevolutionary information (Figure 4), and on the Ube4b system, whose objective is a non-DCA convolutional network. The efficiency therefore reflects how easily the optimizer can navigate the DCA feature landscape, not how the benchmark happens to be scored.

Several limitations frame how these findings should be read. First, every system studied has a deep curated MSA; generalization to shallow-MSA families, where DCA couplings are poorly estimated, will likely require falling back to pLM embeddings despite their higher effective dimensionality. Second, all MD evidence is a structural correlate, not photophysics: quantum yield, excited-state relaxation, and maturation are beyond the scope of ground-state MD, and our single trajectories per variant do not quantify statistical uncertainty in small observables such as SASA shifts. Third, the multi-objective machinery is implemented but untested with competing objectives. Most importantly, all validation here is computational; synthesis and spectroscopic characterization of R9 and intermediates would complete the argument.

The broader point is that surrogate-guided protein design admits analysis, not only benchmarking. When a feature set’s inductive bias can be written as an algebraic property of the representation, and its convergence quantitatively predicted through kernel theory, the choice of representation becomes something one can reason about rather than only measure. Making that shift concrete for low-throughput experimental pipelines is the direction we see for data-efficient protein design.

## 4 Methods

### 4.1 The ALSEBO Framework

ALSEBO is a modular framework that couples generative protein sequence spaces with uncertainty-aware adaptive optimization. The workflow proceeds in four stages (Figure 1): construction and filtering of a candidate sequence library, transformation of each sequence into a numerical feature representation, initialization of a diverse training set, and an iterative Bayesian optimization loop driven by active learning. Each stage is model-agnostic and accommodates user-defined sequence generators, feature representations, and objective functions.

#### 4.1.1 Generative Sequence Space Construction

ALSEBO operates on a user-defined candidate sequence space. In this study, candidate sequences were generated from a latent generative landscape constructed using a variational autoencoder (LGL-VAE). However, the framework does not depend on a specific generative architecture. Any collection of sequences derived from generative latent models, diffusion models, protein language models, directed evolution libraries, or rational design strategies can serve as input. Generated sequences are optionally filtered according to user-defined constraints (e.g., sequence length, mutational distance, or structural compatibility) prior to optimization.

#### 4.1.2 Variational autoencoder architecture

We used a VAE to learn a low-dimensional latent representation of aligned sequences and to generate new, family-consistent samples. The VAE specifies a generative model with **z** and parameters *θ*, consisting of a prior *p_θ_*(**z**) and a conditional likelihood (decoder) *p_θ_*(**x** | **z**). The marginal likelihood of an observed sequence representation **x** is:

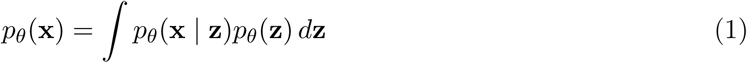

Because the true posterior *p_θ_*(**z** | **x**) is intractable for neural decoders, we introduce a variational approximation *q_ϕ_*(**z** | **x**) (encoder) with parameters *ϕ*:

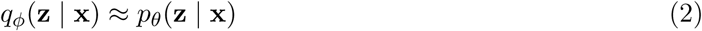

Training maximizes the evidence lower bound (ELBO), obtained by decomposing the log marginal likelihood into a Kullback-Leibler (KL) divergence term and a variational bound:

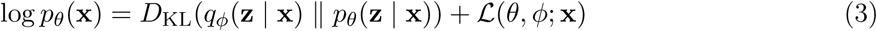

Equivalently, we minimize the negative ELBO, comprising a reconstruction term and a KL regularizer:

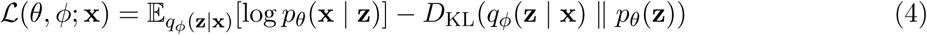

We used a Gaussian latent distribution with a standard normal prior *p_θ_*(**z**) = N(**0**, **I**). The encoder outputs the mean ***µ***(**x**) and diagonal standard deviation ***σ***(**x**), and latent samples are obtained using the reparameterization trick:

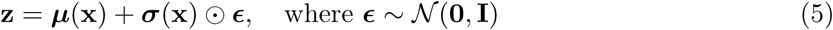

This formulation enables end-to-end optimization by backpropagation through stochastic sampling.

#### Data Representation and Decoding

Aligned amino acid sequences of length *L* were represented as one-hot encoded matrices **X** ∈ {0, 1}*^L^*^×*Q*^, where *Q* = 20 corresponds to the 20 amino acids; each column contains a single 1 indicating the state at that position (gaps, if present in the alignment, were retained as described in the alignment preprocessing). The decoder maps **z** to per-position categorical distributions over the *Q* amino acids via a position-wise Softmax:

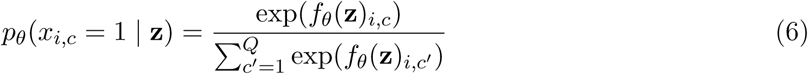

where *f_θ_*(**z**) is the output of the decoder network before the activation function.

#### Hyper-parameters and Training Procedure

The VAE consisted of fully connected encoder/decoder networks with two hidden layers each. Let *L* be the aligned sequence length after preprocessing. The encoder used hidden widths 2*L* followed by *L* units and produced the latent mean and log-variance vectors. The decoder mirrored this structure with hidden widths *L* followed by 2*L* units before the final *L* × *Q* softmax output. ReLU activations were used throughout hidden layers. Models were trained using the Adam optimizer with learning rate 1 × 10^−4^. We applied *L*_2_ weight regularization (penalty 1 × 10^−4^) to mitigate overfitting. Training employed early stopping: optimization was terminated if the reconstruction component of the loss did not improve for 50 consecutive epochs. Training was performed on compute nodes equipped with NVIDIA A100 GPUs.

#### Landscape Generation with Coevolutionary Scoring

To assess evolutionary plausibility across the VAE manifold, we trained a DCA model on the same aligned sequences used for VAE training. DCA defines a Boltzmann distribution over sequences **x**:

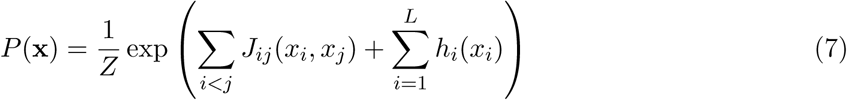

where *J_ij_* are pairwise couplings, *h_i_* are single-site fields, and *Z* is the partition function. From this model, we computed the (unnormalized) Hamiltonian score *H*(**x**):

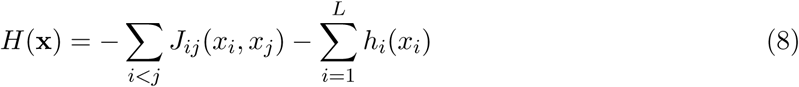

where *i, j* index the sequence positions and *x_i_* represents the discrete state at position *i*.

We generate a LGL by sampling a uniform grid of coordinates **z** spanning the 2D latent space, decoding each coordinate to a categorical distribution (Eq. 6), and then scoring each decoded sequence with the DCA Hamiltonian (Eq. 8). This produced a scalar field over latent space that reports how well decoded sequences conform to the coevolutionary constraints learned from the family alignment.

#### Effective-score interpretation of the LGL

The LGL admits a useful effective-score interpretation that combines the VAE prior with the DCA statistical model. The VAE specifies a standard-normal prior *π_θ_*(**z**) = N(**0**, **I**) and a decoder *p_θ_*(**x** | **z**) over sequences, while DCA defines a Boltzmann distribution over the same sequence space with Hamiltonian *H*_DCA_(**x**). Coupling the two yields the composite score

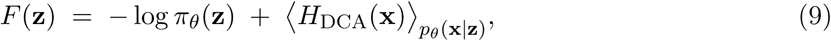

in which the first term is a Gaussian confinement on latent space and the second is the mean coevolutionary penalty of sequences decoded near **z**. This is an effective score rather than a thermodynamically commensurate free energy: the two terms carry different implicit temperatures, and the Hamiltonian dominates the dynamic range by ∼67× (Δ ≈ 2420 vs. 36; SI Figure 5). *F* (**z**) therefore acts as a Hamiltonian-ranked landscape with the VAE prior as a soft domain regularizer: the prior sets *where* the decoder is trustworthy and the Hamiltonian ranks sequences *within* that region. We evaluate ⟨*H*_DCA_⟩ at the MAP-decoded sequence on each grid point, exact in the sharply-peaked-decoder limit.

#### 4.1.3 Sequence Featurization

To enable surrogate modeling, protein sequences are transformed into numerical feature representations. Beyond conventional one-hot encoding, ALSEBO supports three alternative feature modalities: Latent coordinates, DCA features and pLMs Embeddings.^17,34,35^ Sequences can be represented as latent coordinates derived from the trained generative model. These coordinates provide a continuous, low-dimensional embedding that reflects structural and evolutionary constraints learned during model training. The latent coordinates *z*_0_ and *z*_1_ of each generated sequence are retained alongside the sequence for downstream surrogate modeling. We generate a compact, sequence-specific representation using DCA encoding, which projects each aligned sequence into a low-dimensional numeric form derived from an explicit epistatic (Potts) model. As discussed in the previous section, DCA infers local fields (*h_i_*) and pairwise couplings (*J_ij_*) that capture single-site preferences and residue-residue coevolution, respectively. Using these parameters, we define an encoding function (*f*) that assigns each residue (*x_i_*) a single scalar value based on its field term and its couplings to all other positions:

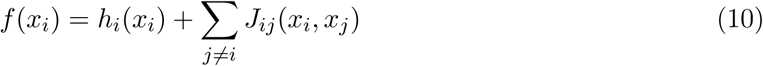

Concatenating *f* (*x_i_*) across positions yields a numerical vector for each sequence, enabling downstream learning while directly incorporating covariation and epistatic constraints into the representation.

This featurization carries an important inductive bias for surrogate-model learning, which can be made explicit by summing Eq. 10 across positions:

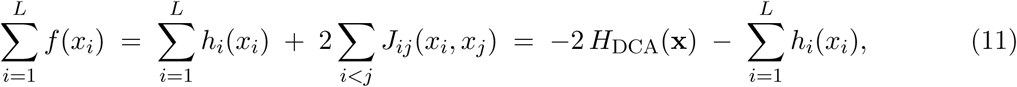

where the last equality substitutes Eq. 8. The DCA feature vector thus contains the negative DCA Hamiltonian as an explicit linear functional: a single direction in feature space (modulo the single-site field correction, which is small relative to the coupling sum for deeply sampled protein families) carries the coevolutionary signal, while the remaining *L* − 1 dimensions provide a *site-resolved decomposition* of that signal. This makes the coevolutionary Hamiltonian linearly accessible to any surrogate that operates on the DCA feature space, in contrast to feature spaces that carry the same information in implicit or nonlinearly distributed form. Whether a specific GP kernel actually exploits this structure depends on kernel choice, hyperparameter fitting, noise, and sample size, and the identity by itself does not prove convergence-rate superiority in a given finite-sample, batched BO setting.

Embeddings derived from pretrained transformer-based models provide contextualized residue-level representations aggregated to sequence-level vectors. Within the ALSEBO framework we extract embeddings from the ESM-2 family, defaulting to the 150M-parameter esm2_t30_150M_UR50D checkpoint for computational efficiency.^17^ The selected featurization strategy is applied to the entire candidate sequence space, yielding a structured feature matrix that serves as input to the downstream surrogate model.

#### 4.1.4 Initialization via Diversity-Aware Sampling

Efficient active learning requires an informative and diverse initial training set. To ensure broad coverage of the candidate space under limited experimental budget, ALSEBO performs diversity-aware initialization using *k*-means clustering in feature space ^36^. Representative sequences nearest to cluster centroids are selected as initial candidates for experimental or computational evaluation; this strategy mitigates early sampling bias and promotes exploration across distinct regions of the generative landscape. Alternatively, previously characterized sequences may be incorporated into the initial training set by projecting them into the same feature space and appending their measured objective values.

#### 4.1.5 Bayesian Optimization and Active Learning Loop

ALSEBO executes an iterative two-step cycle: Gaussian-process regression (GPR) on the accumulated feature-to-objective training set, followed by evaluation of a UCB acquisition function to select the next batch of candidates. Each of *M* user-defined objectives is modeled by its own GPR,

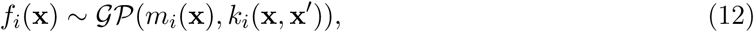

with per-objective mean and kernel *m_i_, k_i_*. Objective-specific acquisition scores

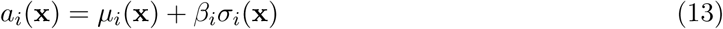

are aggregated into a single acquisition metric by linear scalarization,

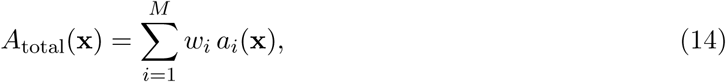

with user-defined weights *w_i_* (uniform by default). The top-*k* sequences by *A*_total_ form the next batch; minimization objectives are handled by negating *f* (**x**). Batches are added to the training set, surrogates are retrained, and the cycle continues until convergence or budget exhaustion. Linear scalarization cannot recover non-convex Pareto fronts; multi-objective extensions using Pareto expected improvement, Thompson sampling, or diversity-aware batch selection are deferred to future work. In this study, *M* = 1 (log-fluorescence fitness).

#### 4.1.6 Construction of the Virtual Objective function (The Digital Twin)

Standard protein-fitness benchmarks such as ProteinGym and FLIP^37,38^ evaluate supervised prediction on fixed train/test splits and are not designed for sequential optimization. To instead benchmark ALSEBO across thousands of optimization trajectories without the resource constraints of in vitro synthesis or explicit-solvent MD, we constructed a virtual objective function (VOF) to serve as a computational surrogate for experiments. We used the comprehensive empirical fitness landscape dataset published by Sarkisyan et al. (2016), which contains experimental fluorescence data for over 32,000 distinct *Aequorea victoria* GFP (avGFP) genotypes. ^28^ This dataset provides a highly rugged benchmarking landscape due to its inclusion of diverse wild-type variants, single-point mutations, and complex multi-point mutations. Following established methodologies from Olivares-Gil et al., we trained a Support Vector Machine (SVM) regressor using DCA features to map sequence variations to experimental fluorescence^39^.

The digital twin achieves a held-out rank correlation (Spearman *ρ*) of 0.75, capturing the dominant fitness trends. Because the SVR is trained on DCA features, the same representation as our primary GPR surrogate, the benchmark that follows may share a basis between oracle and optimizer; we control for this directly with representation-neutral oracles built from one-hot and physicochemical features (Figure 4).

#### 4.1.7 Benchmarking the ALSEBO Framework

We benchmarked ALSEBO against the digital twin using a batch size of 5 candidates per cycle (matching low-throughput cloning-and-characterization throughput) and took the total evaluations needed to reach the oracle optimum as the figure of merit. To probe sensitivity to initial-data scarcity, we ran DCA-featurized campaigns initialized with *T* = 5, 10, and 15 diverse *k*-means-centroid sequences. To probe sensitivity to feature representation, we compared 235-dimensional DCA features (Eq. 10), 640-dimensional ESM-2 embeddings (esm2_t30_150M_UR50D),^17^ and raw 2D LGL latent coordinates under identical conditions, running 100 independent campaigns per representation starting from random individual sequences and recording success rate and mean evaluations to the optimum against a random-search baseline. Because the digital twin objective function and the GPR surrogate were both expressed in DCA coordinates, we controlled for this shared representational basis by reconstructing the oracle using two feature representations that do not encode coevolutionary information: one-hot sequence encodings and physicochemical residue descriptors. We then repeated the complete benchmark against each oracle, using the same starting sequences as the DCA-SVM campaigns (Figure 4B). Finally, to test transfer beyond avGFP, we applied the same protocol to the GFP orthologs cgreGFP and amacGFP (DCA-SVM oracles) ^29^ and to the non-GFP target Ube4b, whose objective was a sequence–fitness convolutional network^33^ trained on deep mutational scanning data, ^32^ reporting convergence rate and mean evaluations to the predicted optimum for each system.

#### 4.1.8 Simulation Setup and Analysis

Starting from the avGFP crystal structure (PDB 2Y0G),^40^ each variant (R6, R7, R8, R9, R10, and the S63T baseline W2) was generated in silico with PyMOL ^41^ and parameterized with the Amber ff14SB force field ^42^ supplemented by chromophore-specific parameters^43^ via the tleap module of AmberTools. ^44^ Systems were solvated in explicit water, neutralized with counter-ions, energy-minimized, and equilibrated before 1 *µ*s production runs with OpenMM v7.7.0^45^ under periodic boundary conditions at 300 K (Langevin thermostat, Particle Mesh Ewald electrostatics). From the aligned trajectories we computed: C*α* RMSF (structural flexibility), SASA (global compaction), chromophore–water RDF (hydration-shell structure), residue-resolved chromophore contact frequencies (*β*-barrel integrity), and the *ϕ*/*τ* dihedral free-energy surfaces of the chromophore (planarity).

For the Ube4b system, the wild-type, the ALSEBO-optimized sequence, and the two nearest natural neighbours in latent space (UniProt F2PJN3 and A0A168CXE7) were modeled from AlphaFold-predicted structures^46^ and parameterized with the AMBER14 all-atom force field. Following the same solvation, neutralization, minimization, and equilibration protocol as above, each system was run for 1.2 *µ*s under identical thermostat and electrostatics settings. From the aligned trajectories we computed C*α* RMSF and performed structural alignment of the optimized model against representative clustered structures from the wild-type dynamics, focusing on the U-box domain and the extended N-terminal region.

## Data and Code Availability

The ALSEBO framework, including the latent generative landscape (LGL-VAE) construction, DCA featurization, and the Bayesian-optimization/active-learning loop, is openly available at https://github.com/dulithaprasanna/ALSEBO and documentation is available at https://dulithaprasanna.github.io/ALSEBO/

All experimental datasets used to build the virtual objective functions are publicly available from their original publications. The avGFP fluorescence fitness data were obtained from Sarkisyan et al. ^28^; the cgreGFP and amacGFP ortholog mutational datasets from González-Somermeyer et al. ^29^; and the Ube4b deep mutational scanning data from Starita et al.,^32^ with the sequence–function convolutional-network oracle adopted from Gelman et al. ^33^. The avGFP crystal structure (PDB 2Y0G) ^40^ was retrieved from the Protein Data Bank, and ESM-2 embeddings were generated with the publicly released esm2_t30_150M_UR50D model. ^17^ Multiple sequence alignments, generated candidate libraries, trained model weights, and molecular-dynamics input files supporting the findings of this study are provided in the repository or are available from the corresponding author upon reasonable request.

## Supporting Information

Supporting Information includes: acquisition landscape evolution across active-learning cycles (SI Figure 1), chromophore dihedral free-energy landscapes for all optimized variants (SI Figure 2), solvent-accessible surface-area distributions from 1 *µ*s MD simulations (SI Figure 3), the perposition mutation-frequency heatmap across BO cycles for the *T* = 10 campaign (SI Figure 4), the three-panel decomposition of the effective-score landscape into the VAE prior, Hamiltonian, and total *F* (*z*) (SI Figure 5), the quantitative GP-UCB information-gain (*γ_T_*) analysis across feature representations (SI Figure 6), fitness-distance correlation (FDC) plots across objective functions and representations (SI Figure 7), additional MD analyses detailing root-mean-square fluctuation (RMSF) and chromophore-neighboring residue interactions (SI Figure 8), and the mutational spread alongside fitness distributions for the evaluated sequence libraries (SI Figure 9).

