## Supporting Information for "Coevolution-informed Bayesian optimization for sample-efficient protein design"

### Supplementary Methods: Fitness Landscape Characterization

To investigate how different sequence representations influence the geometry of the optimization landscape, we characterized the relationship between representation space and the virtual fitness objective using three complementary metrics: fitness-distance correlation (FDC), local smoothness, and Gaussian Process Regression (GPR) predictability. All analyses were performed independently for the DCA, ESM-2, and latent-coordinate representations using the same sequence library and objective values.

#### Fitness-Distance Correlation (FDC)

Fitness-distance correlation quantifies the extent to which fitness varies as a function of distance from the global optimum in a given representation space. For a sequence $i$ represented by feature vector $\mathbf{x}_i$, the Euclidean distance to the highest-fitness sequence $\mathbf{x}^*$ was calculated as:

$$d_i = \|\mathbf{x}_i - \mathbf{x}^*\|_2 \tag{S1}$$

The fitness-distance correlation was then computed as the Pearson correlation coefficient between sequence fitness values $f_i$ and their distances $d_i$:

$$\mathrm{FDC} = \mathrm{corr}(f_i, d_i) \tag{S2}$$

Strong negative FDC values indicate a funnel-like landscape in which fitness decreases monotonically with increasing distance from the optimum, whereas values near zero indicate weak geometric organization and a more rugged landscape.

*

### Local Smoothness

Local smoothness measures the degree to which neighboring sequences exhibit similar fitness values. For each sequence, a $k$-nearest-neighbor graph ($k = 10$) was constructed in the corresponding representation space. The local smoothness metric was defined as the mean absolute fitness difference between each sequence and its neighbors:

$$S = \frac{1}{Nk} \sum_{i=1}^{N} \sum_{j \in \mathcal{N}_k(i)} |f_i - f_j| \tag{S3}$$

where $\mathcal{N}_k(i)$ denotes the set of $k$ nearest neighbors of sequence $i$. Lower values of $S$ correspond to smoother landscapes, where nearby sequences tend to have similar fitness values, whereas larger values indicate greater local epistatic ruggedness.

### GPR Predictability

To assess how readily each representation supports surrogate modeling, Gaussian Process Regression (GPR) models were trained to predict fitness values from sequence features. For each representation, predictive performance was evaluated using five-fold cross-validation. The coefficient of determination ($R^2$) was used as the primary measure of predictability:

$$R^2 = 1 - \frac{\sum_i (f_i - \hat{f}_i)^2}{\sum_i (f_i - \bar{f})^2} \tag{S4}$$

where $f_i$ is the true fitness, $\hat{f}_i$ is the GPR prediction, and $\bar{f}$ is the mean fitness. Higher $R^2$ values indicate that the representation captures fitness-relevant structure that can be efficiently learned by a surrogate model. Because Bayesian optimization relies on accurate surrogate predictions to guide exploration, GPR predictability provides a direct measure of how favorable a representation is for active learning.

### Interpretation

Together, these metrics characterize complementary aspects of the optimization landscape. FDC evaluates global fitness organization relative to the optimum, local smoothness quantifies neighborhood-level ruggedness, and GPR predictability measures the extent to which fitness can be learned from the representation. Representations exhibiting strong negative FDC, low local smoothness values, and high GPR predictive accuracy are expected to provide more favorable landscapes for active machine learning.

### Supplementary Figures

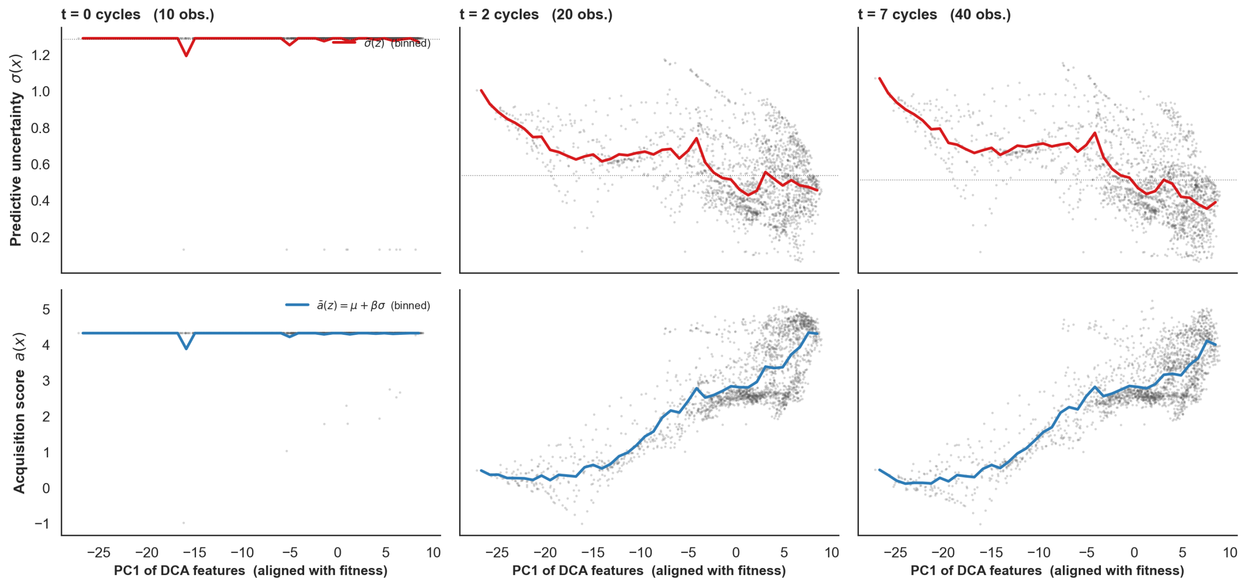


Figure S1: **Evolution of the ALSEBO acquisition landscape.** Predictive uncertainty $\sigma(x)$ (top row) and UCB acquisition score $a(x) = \mu + \beta\sigma$ (bottom row, $\beta = 2$) for the GPR surrogate at $t = 0$, 2, and 7 BO cycles of the $T = 10$ campaign, with all 2,143 library sequences projected onto PC1 of the DCA feature space (aligned with SVR fitness). At $t = 0$ (initial training only) $\sigma$ is uniformly high and $a$ is flat. By $t = 2$, $\sigma$ collapses on the high-PC1 side as the surrogate resolves local fitness structure, and $a$ develops a clear rightward slope. By $t = 7$, $\sigma$ is sharply reduced across explored regions and $a$ concentrates on the high-fitness end, signalling the transition from exploration to exploitation.

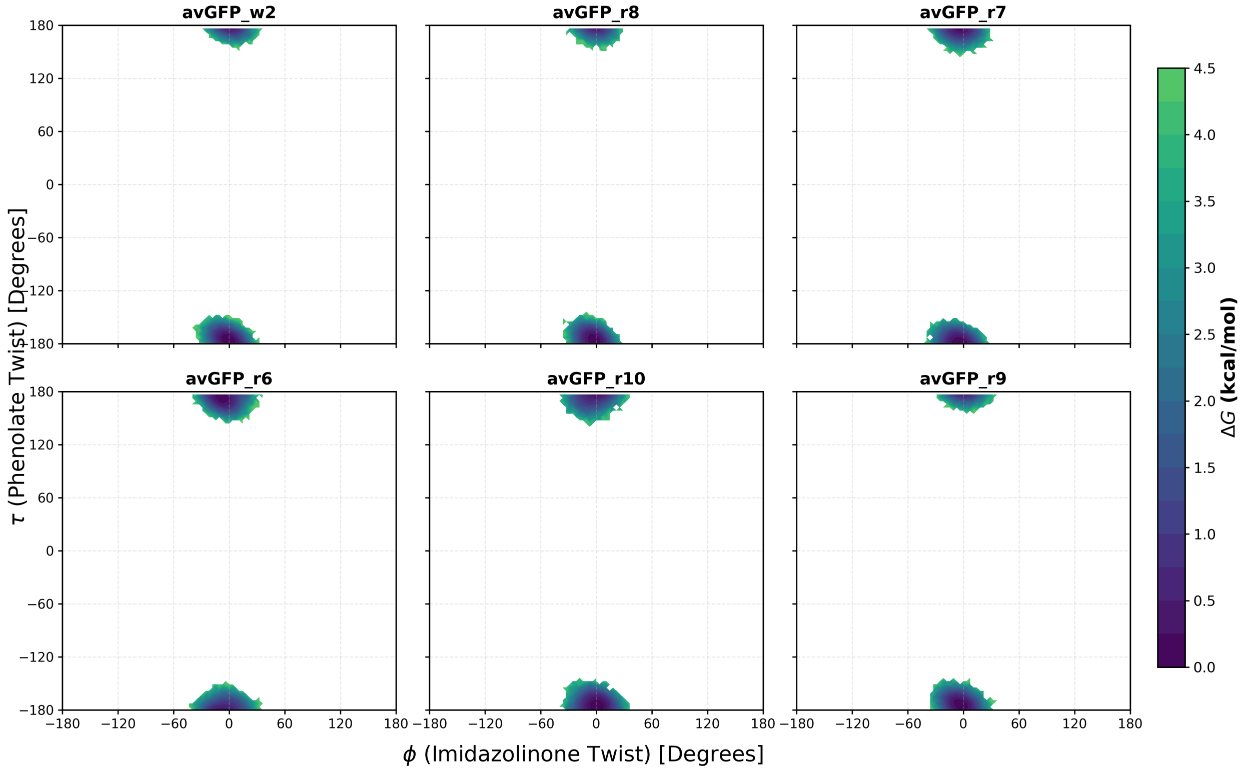


Figure S2: **Chromophore dihedral free-energy landscapes.** The optimized variants (R6–R10) maintain the rigid, coplanar geometries required for extended $\pi$-orbital conjugation, consistent with the restricted torsional energy wells of the functional W2 baseline.

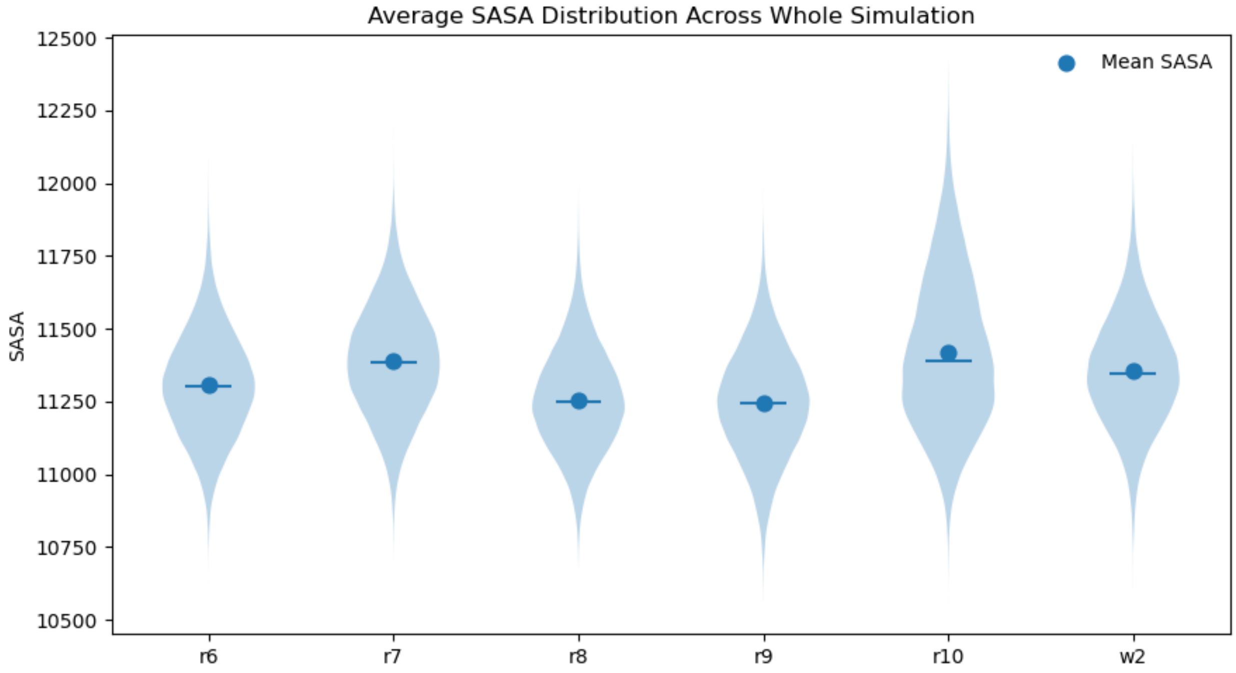


Figure S3: **Structural compaction across 1 $\mu$s simulations.** Solvent-accessible surface-area (SASA) distributions indicate tighter structural compaction for the optimal R9 variant relative to the W2 baseline, consistent with the $\beta$-barrel rigidification in Figure 5D,E of the main text.

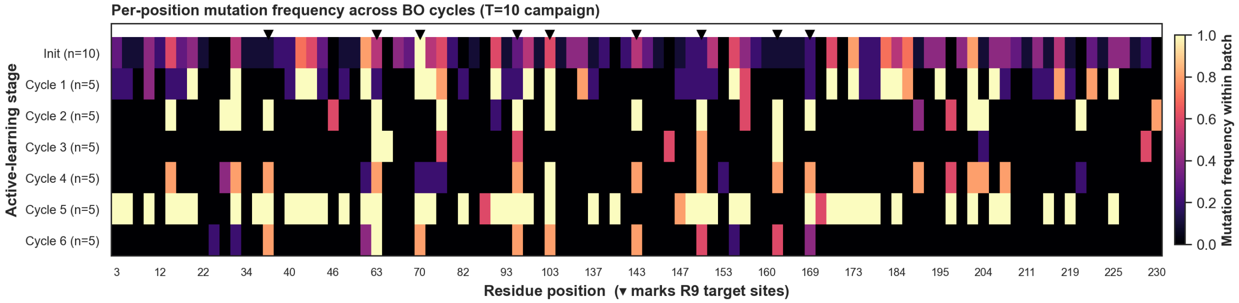


Figure S4: **Per-position mutation frequency across BO cycles.** For the $T = 10$ DCA campaign, each row shows, for one active-learning stage, the fraction of sequences in that batch that carry a mutation at each residue position relative to wild-type avGFP. Only residues mutated at least once across the trajectory are shown ($n = 97$); the nine R9 target positions are marked with ▼ on the top edge. The optimizer explores broad regions of the sequence during early cycles (many mutations at low frequency), with the R9 target sites (e.g., positions 63, 70, 160, 169) rising to high frequency by mid-campaign. This substantiates in a visible form the text's claim that ALSEBO identifies the chromophore-environment substitutions (S65T and neighboring positions) within a few cycles.

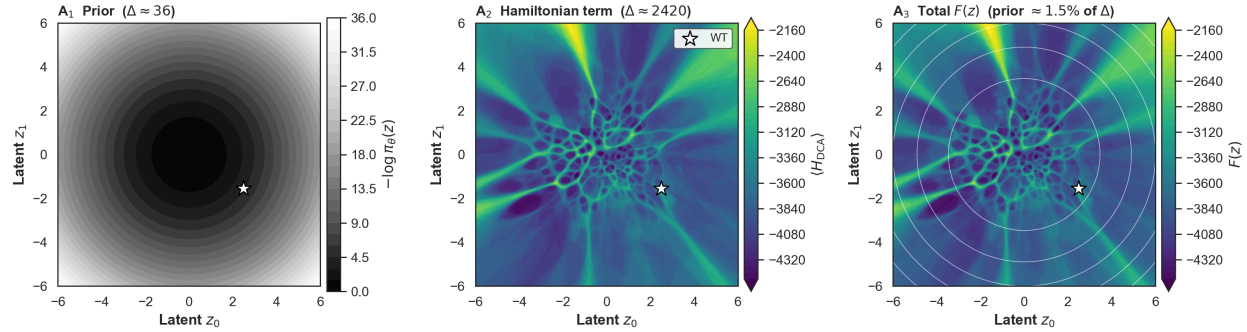


Figure S5: **Decomposition of the effective-score landscape $F(z)$.** ($A_1$) The VAE prior contribution $-\log \pi_\theta(z) = \frac{1}{2}\|z\|^2$ over the full $[-6, 6]^2$ latent domain; its dynamic range is $\Delta \approx 36$. ($A_2$) The DCA Hamiltonian term $\langle H_{\mathrm{DCA}}(x)\rangle$ evaluated at the MAP-decoded sequence on the same grid; its dynamic range is $\Delta \approx 2420$. ($A_3$) The total effective score $F(z) = -\log \pi_\theta(z) + \langle H_{\mathrm{DCA}}\rangle$. $A_2$ and $A_3$ share a common colormap range so the reader directly sees that the Hamiltonian dominates in natural units: the prior contributes only $\sim$1.5% of the dynamic range of $F(z)$. The prior therefore functions as a soft domain regularizer delimiting where the decoder is reliable rather than imposing a per-sequence penalty; its iso-contours are overlaid as thin white lines in $A_3$ for visual reference. The wild-type GFP (white star) sits in a prominent low-$F$ basin.

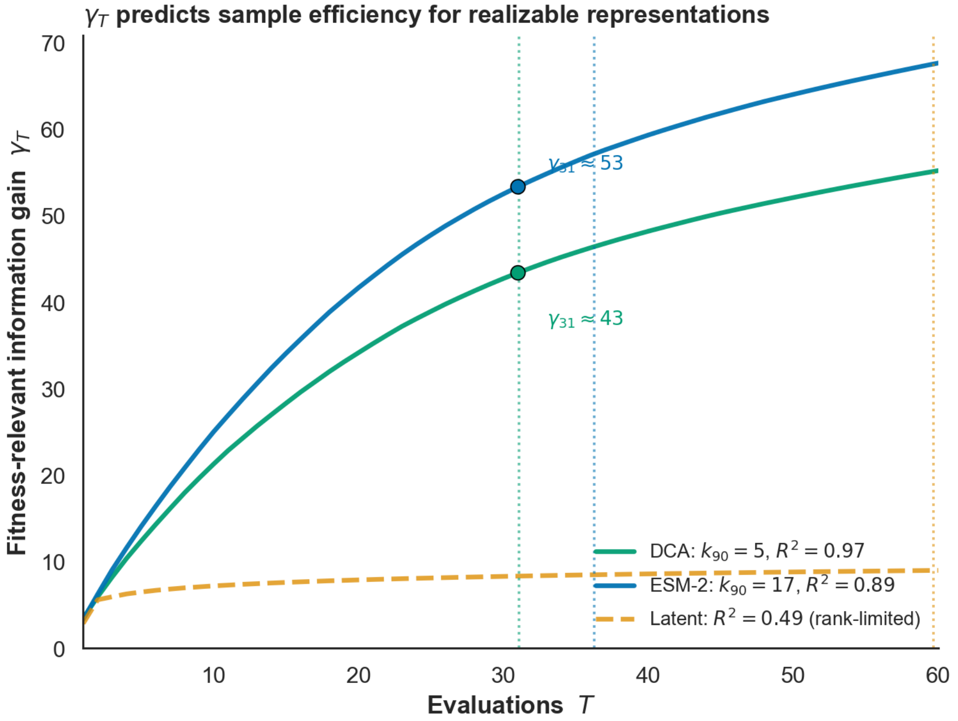


Figure S6: **Fitness-relevant information gain predicts sample efficiency.** Maximum information gain $\gamma_T$ (in the sense of Srinivas et al.) estimated for each feature representation as a function of the number of evaluations $T$, computed by greedy uncertainty sampling under a linear kernel on PCA features weighted by their correlation with the SVR-predicted fitness, so that only fitness-relevant directions contribute (unit signal variance; common observation noise $\sigma_n^2 = 10^{-2}$). Two representations make fitness linearly realizable: DCA (cross-validated $R^2 = 0.97$; $k_{90} = 5$ principal components) and ESM-2 ($R^2 = 0.89$; $k_{90} = 17$). Of these, DCA carries the smaller information gain ($\gamma_{31} \approx 43$ versus 53 at the common horizon $T = 31$). Through the GP-UCB regret bound $R_T = \mathcal{O}(\sqrt{T\gamma_T \log T})$ this predicts the faster convergence observed in the 100-run benchmark (dotted verticals mark the mean steps to optimum, 31 versus 36; main-text Figure 4D). Raw latent coordinates have the smallest $\gamma_T$ yet converge slowest: fitness is not representable in the two-dimensional latent space ($R^2 = 0.49$), a distinct approximation-error regime in which the bound's realizability assumption does not hold.

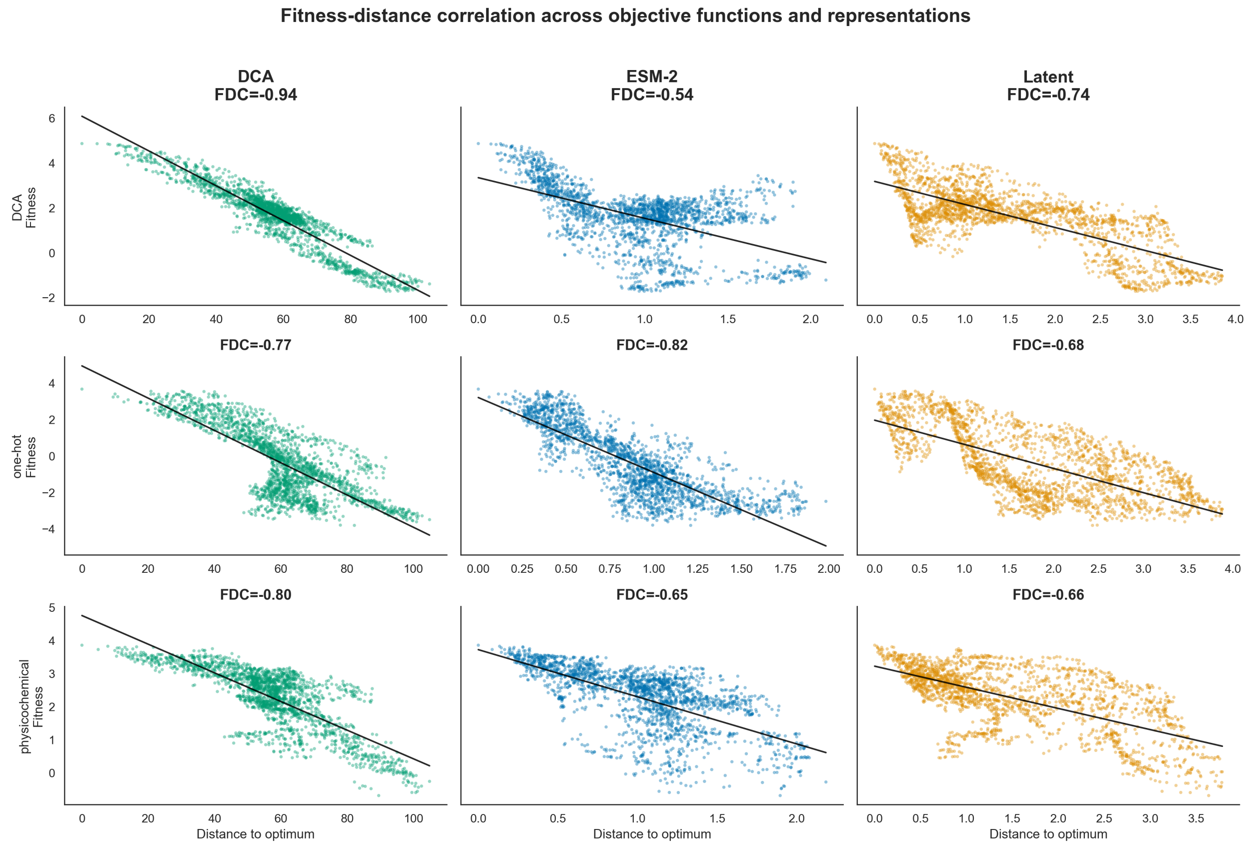


Figure S7: **Fitness-distance correlation (FDC) across different feature representations and objective functions.** Scatter plots illustrating the relationship between sequence fitness and the distance to the global optimum. Columns represent the three evaluated feature spaces (DCA, ESM-2, and Latent coordinates), while rows represent the underlying objective functions used to calculate fitness (DCA, one-hot, and physicochemical). Stronger negative FDC values indicate a more funnel-like, easily optimizable landscape topology.

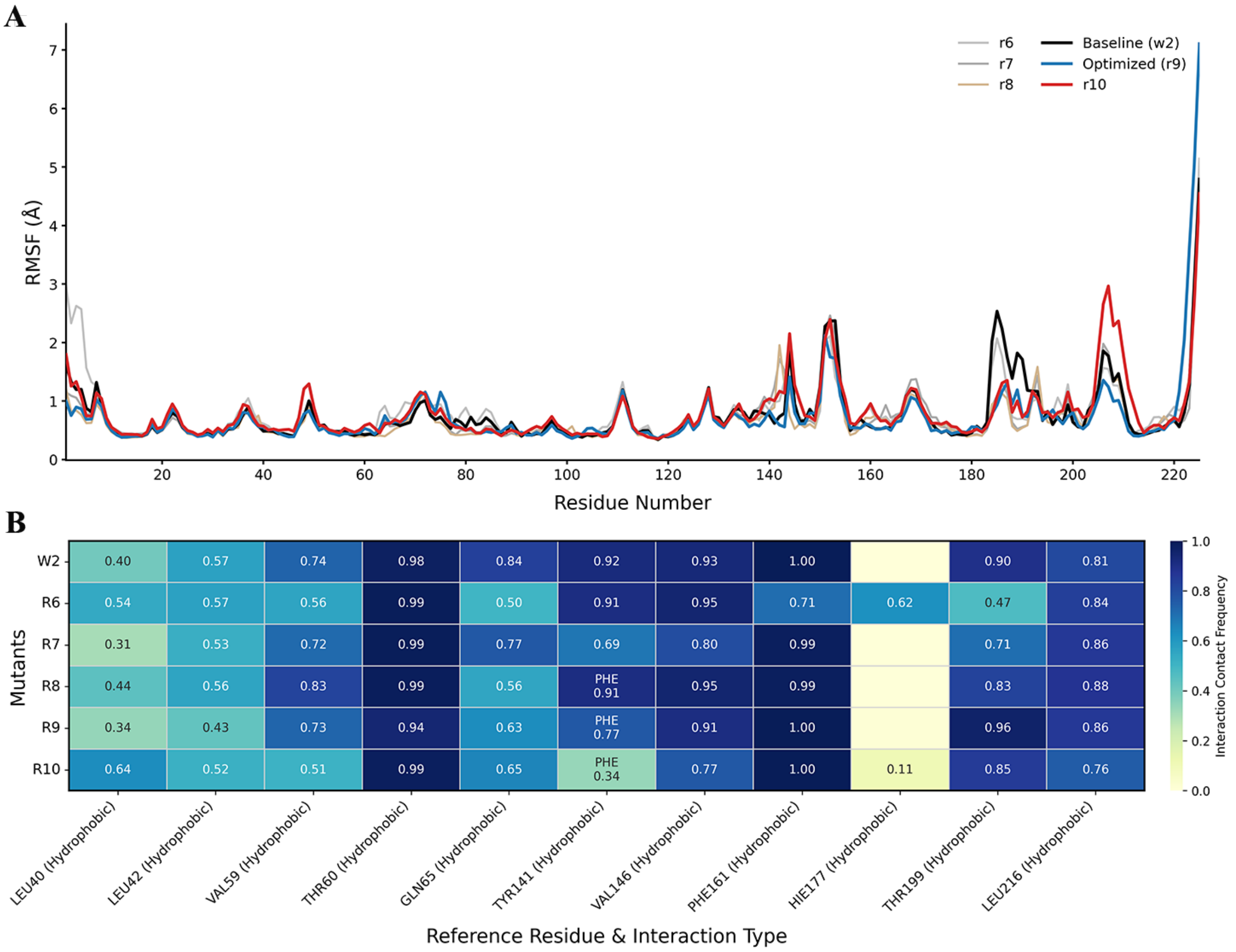


Figure S8: **Molecular dynamics validation of ALSEBO-optimized variants reveals the structural mechanisms of enhanced fluorescence.** A) Root Mean Square Fluctuation (RMSF). B) Chromophore-residue interaction contact frequencies. R9 strictly preserves the essential hydrophobic packing core (e.g., PHE161, VAL146) and successfully accommodates the sfGFP-associated Y→F mutation (position 141) without fracturing the pocket, in contrast to the distorted internal geometries of R6 and R10.

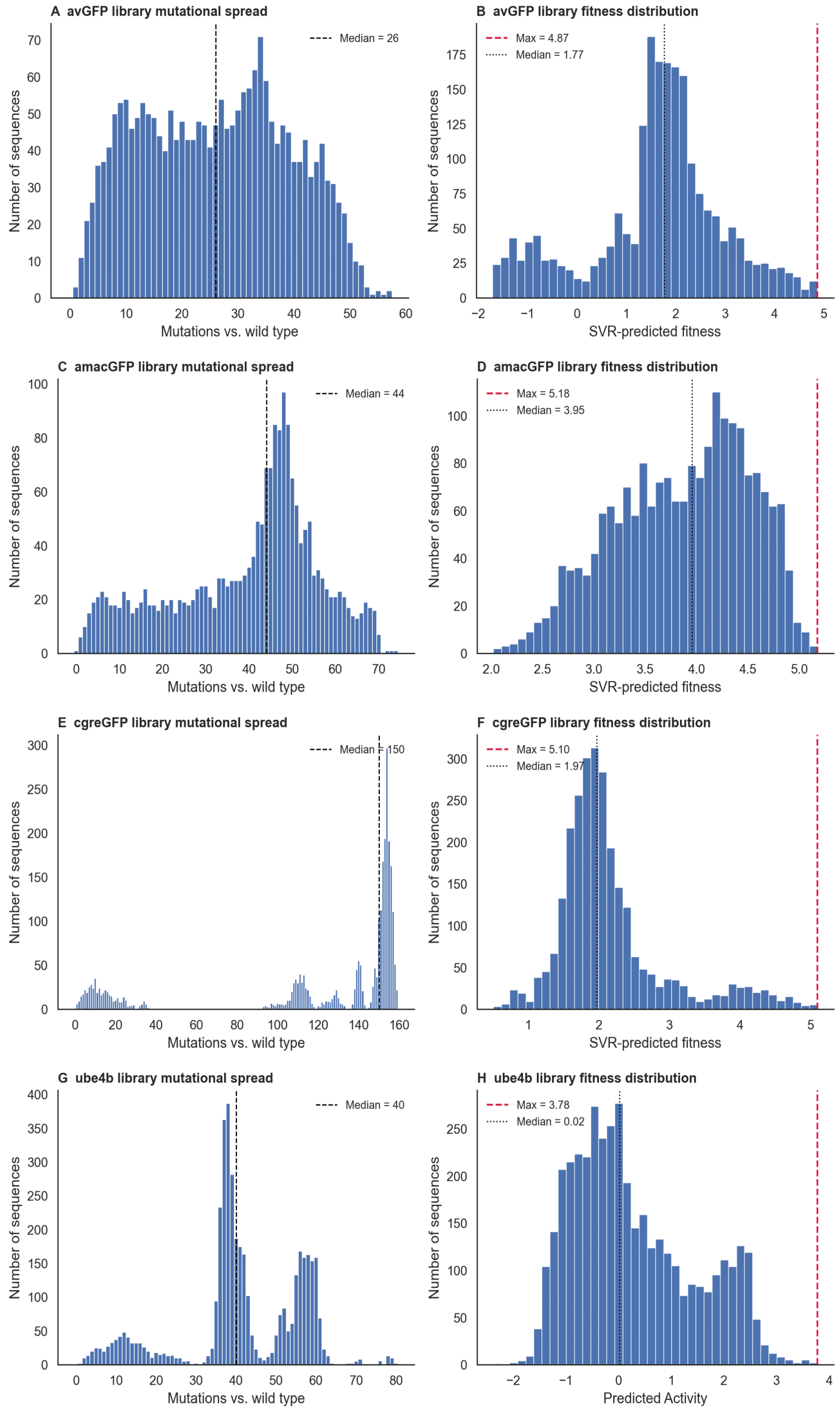


Figure S9: **Mutational spread and fitness distributions for the evaluated sequence libraries.** Left column (A, C, E, G): Distribution of the number of mutations relative to the wild-type sequence for the avGFP, amacGFP, cgreGFP, and ube4b datasets, respectively. Black dashed lines indicate the median mutational distance. Right column (B, D, F, H): Corresponding distributions of the SVR-predicted fitness (or predicted activity) for each generated sequence library. Black dotted lines denote the median fitness of the library, while red dashed lines highlight the global optimum (maximum predicted fitness) within each landscape.
